# A well conserved archaeal B-family polymerase functions as a mismatch and lesion extender

**DOI:** 10.1101/2021.09.10.459716

**Authors:** Xu Feng, Baochang Zhang, Zhe Gao, Ruyi Xu, Xiaotong Liu, Sonoko Ishino, Mingxia Feng, Yulong Shen, Yoshizumi Ishino, Qunxin She

## Abstract

B-family DNA polymerases (PolBs) of different groups are widespread in Archaea and different PolBs often coexist in the same organism. Many of these PolB enzymes remain to be investigated. One of the main groups that are poorly characterized is PolB2 whose members occur in many archaea but are predicted as an inactivated form of DNA polymerase. Herein, *Sulfolobus islandicus* DNA polymerase 2 (Dpo2), a PolB2 enzyme was expressed in its native host and purified. Characterization of the purified enzyme revealed that the polymerase harbors a robust nucleotide incorporation activity, but devoid of the 3’-5’ exonuclease activity. Enzyme kinetics analyses showed that Dpo2 replicates undamaged DNA templates with high fidelity, which is consistent with its inefficient nucleotide insertion activity opposite different DNA lesions. Strikingly, the polymerase is highly efficient in extending mismatches and mispaired primer termini once a nucleotide is placed opposite a damaged site. Together, these data suggested Dpo2 functions as a mismatch and lesion extender, representing a novel type of PolB that is primarily involved in DNA damage repair in Archaea. Insights were also gained into the functional adaptation of the motif C in the mismatch extension of the B-family DNA polymerases.

## INTRODUCTION

Cellular organisms code for multiple DNA polymerases that play crucial roles in the chromosome duplication and the genome integrity maintenance during the normal growth and under stressed conditions. Eight different families of DNA polymerases (pols) are known based on their amino acid sequences, and as many as 17 DNA pols are encoded in human [1]. Some polymerases are devoted to chromosome replication (replicase) while others are specialized for DNA damage repair. Bacterial replicases for chromosome replication are of the C-family, and those in the organisms of Eukarya and Archaea belong to the B-family or D-family [2; 3; 4]. Replicative polymerases possess both the polymerase and exonuclease domains and replicate undamaged DNA with high fidelity and processivity. In contrast, most specialized DNA pols are of X- and Y-family. These pols are often devoid of any proof-reading activity and replicate DNA with reduced fidelity and processivity [1]. A noticeable exception of specialized pols is the eukaryotic Pol ζ, a B-family DNA polymerase, which plays important role in the eukaryotic translesion DNA synthesis (TLS) by functioning as a mismatch and lesion extender.

Sulfolobales organisms code for four DNA polymerases such as *Sulfolobus islandicus, Sulfolobus acidocaldarius* and *Saccharolobus solfataricus* P2 (formerly *Sulfolobus solfataricus*). These DNA pols were initially named as Dpo1, Dpo2, Dpo3 and Dpo4/Dbh among which the first three belong to the B family (also known as PolB1, PolB2 and PolB3) whereas the last is a Y-family pol [5; 6; 7]. DNA pols encoded in *S. solfataricus* were characterized in vitro in different research laboratories, and these analyses have revealed the Dpo1 and Dpo3 enzymes are high-fidelity DNA polymerases exhibiting the 3’-5’ exonuclease activity, and this is consistent with their predicted function in the processive DNA replication in this crenarchaeon [8; 9; 10]. The *S. solfataricus* Dpo4 represents the most extensively characterized Y-family DNA pol. This enzyme is capable of bypassing various DNA lesions [11; 12; 13; 14], suggesting it could be responsible for the translesion synthesis in this organism. However, the encoding gene does not show any DNA damage-inducible expression in all tested Sulfolobales organisms, including *S. acidocaldarius, S. solfataricus*, and *S. islandicus* [15; 16; 17; 18; 19], and it does not play a role in the targeted mutagenesis as we have demonstrated with the *S. islandicus* Δ*dpo4* mutant [20]. The only DNA pol gene that does show the damage-inducible expression is *dpo2* coding for a PolB2 enzyme [15; 16], and it has been further shown that *dpo2* is solely responsible for the DNA damage-induced mutagenesis in *S. islandicus* [20]. Nevertheless, members of the PolB2 subfamily were regarded as inactive polymerases since they carry amino acid substitutions at the catalytic center [21]. Consequently, whether Dpo2 could be an active polymerase represents very important question in the TLS study in Dpo2-encoding organisms, and if so, it would be intriguing to know how this unique DNA pol could contribute to DNA damage repair in these archaea.

Herein, we biochemically characterized the *S. islandicus* Dpo2. Recombinant Dpo2 protein was obtained from the native host and investigated for its capability of DNA polymerization, proofreading and lesion bypass. We found that Dpo2 is a robust DNA pol in nucleotide incorporation but devoid of the 3’-5’ exonuclease activity. This unique DNA pol replicates undamaged DNA with a high replication fidelity that is comparable to that of Dpo1, the main replicase of the organism. We further demonstrate that Dpo2 is very efficient in extension of mismatched primer termini as well as primer termini opposite to DNA lesions. Together, these results indicated that the PolB2 enzymes are specialized DNA polymerases that can play very important roles in archaeal DNA damage repair.

## MATERIALS AND METHODS

### *Sulfolobus* strains and growth conditions

*S. islandicus* E233S (Δ*pyrEF*Δ*lacS*) [22], derived from *S. islandicus* REY15A, the wild-type strain [23], was employed as the host for expression of recombinant DNA polymerases including *S. islandicus* Dpo2 and Dpo1, and *S. solfataricus* Dpo2. *Sulfolobus* strains were grown in SCV (0.2% sucrose, 0.2% casamino acids, 1% vitamin solution plus basic salts) or ACV media (0.2% D-arabinose, 0.2% casamino acids, 1% vitamin solution plus basic salts) at 78°C as previously described [24].

### Expression and purification of DNA polymerases from *S. islandicus*

The *S. islandicus* Dpo2-expression plasmid and its native host strain were constructed previously [25]. Dpo1 (SiRe_1451) and SsoDpo2 (Sso1459) encoding genes were amplified by PCR using corresponding oligos listed in Supplementary Table S1 and individually cloned into pSeSD, an arabinose inducible expression vector [24]. Construction of the expression plasmids, expression of the DNA pol genes and purification of the encoded proteins from *S. islandicus* E233S were conducted as previously described [26]. Briefly, 20-500 ng plasmid DNA were used for electroporation transformation for each plasmid, and the colonies appeared on the selective plates (SCV) were check for DNA insert by colony PCR, and the target genes were verified by sequencing of the PCR product. Transformants carrying each expression plasmid were firstly cultured in SCV, the non-induction medium, for cell growth. Then, the cultures were transferred into ACV, the induction medium, for protein expression. Cell mass was harvested from ca. 11 L of ACV cultures and used for the purification of Dpo2 and Dpo1 individually by the following procedures. Cell pellets were resuspended in Buffer A (50 mM Tris-HCl, 200 mM NaCl, 30 mM Imidazole, pH 7.5) supplemented with 1x protease inhibitor cocktails and 10 μg/ml DNase I. Cell lysates were obtained by passing the cell suspension through a high-pressure homogenizer (JNBIO). Cell debris in the lysates were removed by centrifugation at 15000 × g for 40 min and the supernatant was filtered through a 0.45 μm filter. The clarified supernatant was then applied to a Histrap HP column (Cytiva) and the target protein bound to the Ni column via the specific His tag-Ni ion interaction was then eluted with Buffer B (50 mM Tris-HCl, 200 mM NaCl, 500 mM Imidazole, pH 7.5). Further purification of recombinant protein was different for Dpo2 and Dpo1. In the case of Dpo2, pooled fractions were diluted using Buffer C (20mM Tris-HCl, pH 8.0), and applied onto a Heparin HP column (Cytiva). Proteins bound to the Heparin column were then eluted by a linear gradient of buffer D (20 mM Tris-HCl, 1 M NaCl, pH 8.0) over a 25x column volume. Further purification of Dpo1 was conducted by size exclusion chromatography with a Superdex 200 increase 10/300 GL column (Cytiva). Fractions containing each DNA pol of a high purity were pooled and concentrated using a 10K protein concentrator (Millipore). Concentrated proteins were preserved at −20 °C in the presence of 50% glycerol. The concentration of each protein was determined using Bradford assay [27], with BSA of known concentrations as standards.

### DNA substrates

All synthetic oligos including unlabeled, FAM-labeled primers, undamaged templates, and templates containing base modifications were synthesized and purified by HPLC at Genewiz (Suzhou, CN) or Sangon Biotech (Shanghai, CN). The exception was a CPD-containing oligo, which was synthesized and purified by Gene Link (Elmsford, NY, USA). The sequence of primers and templates were listed in Supplementary Table S1. DNA substrates were prepared by annealing corresponding primer stand and template strand at 1:1.5 ratio using a thermal cycler, in which the temperature was decreased by 0.2 °C each cycle for 350 cycles after denaturation at 95 °C for 5 min.

### Optimization of reaction conditions

The optimal pH was determined using Bis-Tris based buffer system in the pH range of 6.0-7.2 and Tris-HCl system in the range of 6.8-8.8. Salt concentrations of NaCl and KCl were screened at the pH 8.0 of Tris-HCl buffer. At optimal pH and salt concentration, the reaction temperature and concentration of metal ions was optimized. From these experiments, the optimal condition was determined for the Dpo2 reaction in the solution containing 50 mM Tris-HCl (pH8.0), 40 mM KCl, 10 mM MgCl_2_, 0.1 mg/mL BSA, at 60 °C, and it was used for the following analyses.

### Primer extension assay

Primer extension was set up in a 10 μl reaction system containing 50 nM substrates, DNA polymerases with indicated concentrations, 100 μM of either all four dNTPS or each dNTP individually, 50 mM Tris-HCl (pH 8.0), 40 mM KCl, 10 mM MgCl_2_, 0.1 mg/mL BSA using undamaged or damaged templates. The assay was carried out at 60 °C for 10 min or with the time periods indicated in each experiment. The reaction was terminated by addition of 10 μl 2x loading dye solution (1x TBE, 8 M Urea, 10 mM EDTA, 0.1% bromophenol blue), followed by denaturation at 95 °C for 5 min and immediate chilling on ice. Replication products were resolved by 18% urea polyacrylamide gel electrophoresis and visualized by an Amersham ImageQuant 800 biomolecular imager (Cytiva).

### Proofreading assay

The assay was performed essentially as described in primer extension assay, except dNTPs were omitted from the reaction.

### Steady-state kinetics analysis

Steady state kinetics was performed as described [28]. To ensure that the reaction was in the linear range, products formation was kept to less than 20% of the starting substrate. For misincorporation and mismatch extension kinetic assay, each 10 μl reaction contained 50 nM 5’ FAM-labelled substrate, 1000 nM corresponding unlabeled cold DNA substrate with the same DNA sequence, 350 nM Dpo2 protein. The reaction was initiated by addition of dNTP with varying concentrations and was terminated by mixing with 10 μl 2x loading dye solution (1x TBE, 8 M urea, 10 mM EDTA and 0.03% bromophenol blue) and heating at 95 °C for 5 min. Products were resolved in a 18% urea-PAGE gel and visualized by an Amersham ImageQuant 800 biomolecular imager. The percentage of products formation was quantitated using ImageQuant software and the velocity of dNTP incorporation was calculated by dividing the yield of products formed by the respective time of the reaction at each concentration of dNTP. The data was fitted into Michaelis-Menton equation using Graphpad prism software, from which the apparent K_cat_ and K_m_ values were determined. The misinsertion frequency was expressed as *f*_inc_ = (K_cat_/K_m_) _incorrect_/(k_cat_/K_m_)_correct_. The intrinsic efficiency of mismatch extension of Dpo2 on mismatch extension was calculated as described [29] using the equation: *f*^0^_ext_ = (K_cat_/K_m_)_mismatch_/(K_cat_/K_m_)_matched_, which measures the relative probability of extending mismatched termini in competition with matched primer, in the limit of zero next nucleotide and has been widely used to evaluate the mispair extension ability of DNA polymerases [29; 30; 31]. The substrates used for misincorporation kinetics including P2-T1, P2-T1-A, P2-T1G-A and P2-T1C as indicated in Supplementary Table S2. The substrates used for mismatch extension kinetics including P3-T1N (N=A, T, G or C), P3T-T1N (N=A, T, G or C), P3G-T1N (N=A, T, G or C) and P3C-T1N (N=A, T, G or C).

## RESULTS

### *S. islandicus* Dpo2 is an active DNA polymerase devoid of the exonuclease activity

In a previous work, we showed that *dpo2* is the only DNA polymerase gene essential for DNA damage-induced mutagenesis in *S. islandicus* [20]. To test how this polymerase could be related to other B-family DNA polymerases, multiple sequence alignments were conducted for the *S. islandicus* Dpo2 protein together with a selected set of B-family polymerases, including those of *Saccharomyces cerevisiae, E. coli* and a few archaea. This analysis revealed that the PolB2 enzymes from different archaeal species possess a relatively conserved polymerase domain, including conserved PolA, PolB and PolC motifs that are normally conserved in other B-family DNA polymerases, albeit its PolC motif carries amino acids substitutions at conserved Y and D (Fig. 1A). In contrast, the exonuclease domain is either absent or much more diverged from those present in the B-family replicative polymerases (Fig. 1A, Fig. S1).

**Figure 1.**
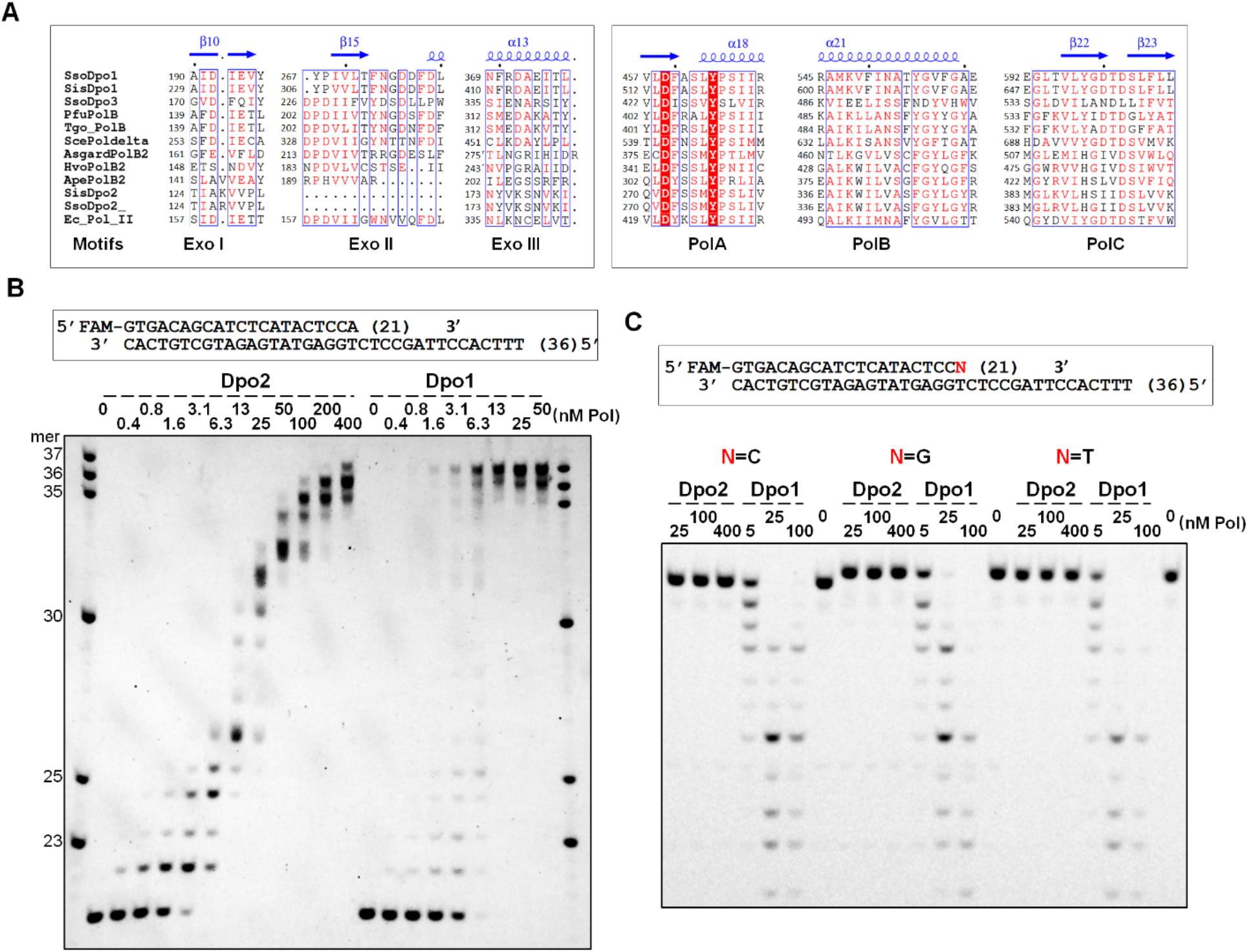
Dpo2 is robust in DNA polymerization but deficient in proofreading activity. (**A**) Sequence alignment of a few selected B-family DNA Polymerases. Only the selected regions of the exonuclease domain and polymerase domain are shown and the full sequence alignment was shown in Fig S1. SsoDpo1, *S. solfataricus* Dpo1. SisDpo1, *S. islandicus* Dpo1. SsoDpo3, *S. solfataricus* Dpo3. PfuPolB, *Pyrococcus furiosus* PolB. Tgo_PolB, *Thermococcus gorgonarius* PolB. ScePoldelta, The catalytic subunit of *Saccharomyces cerevisiae* Pol δ. AsgardPo1B2, Candidatus Thorarchaeota archaeon PolB2. HvoPolB2, *Haloferax volcanii* PolB2. ApePolB2, *Aeropyrum pernix* PolB2. Ec_Pol_II, *E. coli* Pol II. Structures of SsoDpo1 (1S5J) was used as the templates for the structure-based sequence alignment. The secondary structural elements shown above the sequences were retrieved from the structure file of SsoDpo1 (1S5J) (**B**) Primer extension activities of Dpo2 and Dpo1. Reactions were set up with 50 nM primer-template, 100 μM dNTPs and a concentration gradient of Dpo2 or Dpo1 (indicated above their gel images in each panel). After incubation at 60 °C for 10 min, extension products were analyzed by denaturing PAGE. Note: Dpo1 yielded an extension product of 37 nt, which is one nucleotide longer than the template (36 nt), indicative of a strong TdT (terminal transferase) activity of the enzyme. In contrast, Dpo2 only showed a low TdT activity. (**C**) Proofreading by Dpo2 and Dpo1. Exonuclease assay was set up with 50 nM mismatched primer-template and a gradient concentration of Dpo2 or Dpo1 in the absence of dNTPs. After incubation at 60 °C for 5 min, the products were analyzed by denaturing PAGE. N denotes each of the four possible primer terminal nucleotides as indicated.

To characterize Dpo2 biochemically, the encoding gene of *S. islandicus* was expressed in its native host and purified into an apparent homogeneity (Fig. S2). The purified Dpo2 was assayed for the basic properties and for the optima of the primer extension reaction (Fig. S3), using the substrate shown in Fig. 1B. These included determination of its optimal values in reaction pH, temperature and salt content as well as the metal ion preference. As shown in Fig. S3A, the activity of Dpo2 increased along with the increase of pH from 6.0 to 8.0, before shallowing down at pH 8.8. Longest synthesized DNA fragments appeared in the reactions of pH 8.0 and 8.4, suggesting this pH range is optimal for the polymerase. As a result, subsequent optimization of the Dpo2 assay was conducted with buffers containing 50 mM Tris-Cl pH 8.0. The effect of salt on the activity of Dpo2 was tested with both KCl and NaCl. We found that the *S. islandicus* Dpo2 was most active at the low salt buffer, and in fact, 80 mM KCl or 20 mM NaCl could already inhibit the Dpo2 activity (Fig. S3B). Six different divalent metal ions (Mg^2+^, Mn^2+^, Ca^2+^, Zn^2+^, Ni^2+^, Fe^2+^) were tested for their capability of supporting the polymerization activity, and this revealed that both magnesium and manganese ions supported the Dpo2 activity. Furthermore, Dpo2 showed a higher activity in the presence of the equal concentration of the manganese ion relative to the magnesium ion (Fig. S3D), as reported for many specialized DNA polymerases [32; 33; 34; 35; 36]. Nevertheless, Mg^2+^ was used in the following analysis, considering a much higher physiological concentration for this metal ion in different cells. In addition, the optimal reaction temperature and dNTP concentration determined for Dpo2 were 55-65 °C and 100-500 μM, respectively (Fig. S3). To this end, the optimized buffer system for Dpo2 was defined as 50 mM Tris-HCl pH 8.0, 40 mM KCl, 0.1 mg/ml BSA, 10 mM MgCl_2_, 100 μM dNTPs, which was employed for all primer extension assays.

Using the optimized reaction system, we examined the nucleotide incorporation activity of Dpo2, in comparison with Dpo1, the replicase of this crenarchaeon [8]. The enzyme concentrations tested for Dpo2 and Dpo1 were 0.4 - 400 nM and 0.4 - 50 nM, respectively, and this revealed that Dpo2 manifested DNA polymerization at a concentration as low as 3 nM, and the amount of primer consumed by Dpo2 in this assay was comparable to that converted by Dpo1 at the identical or a very similar enzyme concentration (Fig. 1B). These results indicated that Dpo2 exhibits a robust nucleotide incorporation activity. Noticeably, while the replicase readily extended the primer into full length products (with 13 nM Dpo1), the Dpo2 polymerization yielded DNA fragments of different sizes in the reactions with the same or a higher enzyme concentration (Fig. 1B). These data suggested that the *S. islandicus* Dpo2 is a distributive polymerase, relative to the processive Dpo1 enzyme.

To test if the *S. islandicus* Dpo2 could perform proof-reading during DNA synthesis, the mismatched primer-templates (including T:C, T:G and T:T mismatches) were mixed individually with Dpo2 (25, 100 or 400 nM) as well as Dpo1 (5, 25, or 100 nM), the latter of which is known to possess the 3’-5’ exonuclease activity. After incubation at 60 °C for 5 min, samples were analyzed by the denaturing PAGE. As shown in Fig. 1C, while 25 nM Dpo1 effectively degraded all three primers from the 3’-terminus, generating a ladder of degraded oligonucleotides, 16-fold more Dpo2 enzyme did not show any detectable 3’ −5’ exonuclease activity since full-length primers remained intact in the reaction with 400 nM Dpo2.

Taken together, the *S. islandicus* Dpo2 represents a unique PolB exhibiting robust nucleotide-incorporation, poor processivity and no detectable exonuclease activity.

### Dpo2 replicates undamaged DNA with high fidelity

Next, we sought to decipher kinetic parameters of nucleotide incorporation by this unique B-family enzyme, using the steady-state kinetic assay described in the Materials and Methods. Dpo2 was evaluated for the fidelity of nucleotide incorporation opposite each of the four template bases. As summarized in Table 1, Dpo2 is clearly able to discriminate correct and incorrect incoming nucleotide, as it incorporated correct nucleotides opposite different template base with highest efficiency (K_cat_/K_m_) and with lowest K_m_ values. Overall, insertion of a wrong nucleotide (misincorporation) by Dpo2 occurred at a frequency ranging from 2.8 × 10^−5^ (for inserting a C opposite a template base C) to 5.37 × 10^−4^ (for inserting a G opposite an A). Thus, the misincorporation frequency of this unique PolB on four different template base is from 10^−4^ to 10^−5^, falling into the same range of the replication fidelity by the *S. solfataricus* replicase Dpo1 at 37 °C [37] and its exonuclease-minus mutant, Dpo1 exo^−^ at 55 °C [9]. Thus, Dpo2 is a high-fidelity DNA polymerase on undamaged DNA templates.

**Table 1.** Steady-state kinetic parameters of deoxynucleotide incorporation by Dpo2 on undamaged DNA.

| Template<br>base | Incoming<br>dNTP | $K_m$<br>($\mu\text{M}$) | $K_{cat}$<br>($\text{min}^{-1}$) | $K_{cat}/K_m$<br>($\mu\text{M}^{-1} \text{min}^{-1}$) | $f_{inc}$ |
| --- | --- | --- | --- | --- | --- |
| A | A | $1205 \pm 201$ | $0.0112 \pm 0.0022$ | $9.3 \times 10^{-6}$ | $1.91 \times 10^{-4}$ |
| | T | $39.9 \pm 14.9$ | $1.94 \pm 0.41$ | $4.9 \times 10^{-2}$ | 1 |
| | G | $902 \pm 188$ | $0.0235 \pm 0.00076$ | $2.6 \times 10^{-5}$ | $5.37 \times 10^{-4}$ |
| | C | $488 \pm 56.2$ | $0.00587 \pm 0.00015$ | $1.2 \times 10^{-5}$ | $2.48 \times 10^{-4}$ |
| T | A | $118 \pm 10.6$ | $11.7 \pm 5.91$ | $9.9 \times 10^{-2}$ | 1 |
| | T | $1616 \pm 91$ | $0.0234 \pm 0.0021$ | $1.5 \times 10^{-5}$ | $1.46 \times 10^{-4}$ |
| | G | $1454 \pm 131$ | $0.0299 \pm 0.0040$ | $2.1 \times 10^{-5}$ | $2.08 \times 10^{-4}$ |
| | C | $413 \pm 83.6$ | $0.00156 \pm 0.00028$ | $3.8 \times 10^{-6}$ | $3.81 \times 10^{-5}$ |
| G | A | $982 \pm 75.9$ | $0.0118 \pm 0.0020$ | $1.2 \times 10^{-5}$ | $1.17 \times 10^{-4}$ |
| | T | $1773 \pm 107$ | $0.0279 \pm 0.0077$ | $1.6 \times 10^{-5}$ | $1.53 \times 10^{-4}$ |
| | G | $2474 \pm 628$ | $0.0257 \pm 0.0053$ | $1.0 \times 10^{-5}$ | $1.01 \times 10^{-4}$ |
| | C | $40.6 \pm 5.78$ | $4.17 \pm 0.92$ | $1.0 \times 10^{-1}$ | 1 |
| C | A | $1230 \pm 168$ | $0.0241 \pm 0.0047$ | $1.9 \times 10^{-5}$ | $3.17 \times 10^{-4}$ |
| | T | $1428 \pm 130$ | $0.0084 \pm 0.0015$ | $5.9 \times 10^{-6}$ | $1.00 \times 10^{-4}$ |
| | G | $62.1 \pm 8.54$ | $3.63 \pm 0.29$ | $5.8 \times 10^{-2}$ | 1 |
| | C | $684 \pm 231$ | $0.00112 \pm 0.00032$ | $1.6 \times 10^{-6}$ | $2.80 \times 10^{-5}$ |

$K_m$ and $K_{cat}$ values were determined by quantification of gel bands corresponding to substrates and products using ImageQuantTL, and the data were fitted into the Michaelis-Menten equation using Graphpad Prism. The nucleotide misincorporation ratio ( $f_{inc}$ ) was expressed as $(K_{cat}/K_m)_{incorrect}/(K_{cat}/K_m)_{correct}$ . SD values means standard variations from three independent experiments.

### Dpo2 is proficient in extension of mismatched primer termini

To test if Dpo2 could extend mismatched base pair ends, we determined the ability of Dpo2 to elongate 4 matched and 12 mismatched primer-templates (Table S2), using the steady state kinetics assay. As summarized in Table 2, the frequencies of Dpo2 (*f*^0^_ext_) in mismatch extension from A:A, A:G and G:T mispairs were estimated to 4.5 × 10^−1^, 1.3 × 10^−1^ and 2.6 × 10^−1^ respectively, and *f*^0^_ext_ values for extension from most of the rest mispairs were found to be in the order of 10^−2^. These results indicated that Dpo2 can effectively extend mismatched DNA ends.

**Table 2.** Steady-state kinetics parameters for mispair extension by Dpo2.

| Template | Primer | $K_m$ | $K_{cat}$ | $K_{cat}/K_m$ | $f_{ext}^0$ |
| --- | --- | --- | --- | --- | --- |
| base | base | ($\mu M$) | ($\text{min}^{-1}$) | ($\mu M^{-1} \text{min}^{-1}$) | |
| A | A | $55.0 \pm 5.0$ | $1.76 \pm 0.067$ | $3.2 \times 10^{-2}$ | $4.5 \times 10^{-1}$ |
| | T | $55.2 \pm 14.5$ | $3.96 \pm 0.34$ | $7.2 \times 10^{-2}$ | 1 |
| | G | $323 \pm 84.7$ | $3.09 \pm 0.21$ | $9.6 \times 10^{-3}$ | $1.3 \times 10^{-1}$ |
| | C | $1129 \pm 222$ | $1.03 \pm 0.095$ | $9.1 \times 10^{-4}$ | $1.3 \times 10^{-2}$ |
| T | A | $19.1 \pm 2.66$ | $35.9 \pm 3$ | 1.9 | 1 |
| | T | $325 \pm 59.6$ | $1.43 \pm 0.34$ | $4.4 \times 10^{-3}$ | $2.3 \times 10^{-3}$ |
| | G | $187 \pm 22$ | $21.5 \pm 8.91$ | $1.1 \times 10^{-1}$ | $6.1 \times 10^{-2}$ |
| | C | $886 \pm 76.1$ | $19.7 \pm 5.89$ | $2.2 \times 10^{-2}$ | $1.2 \times 10^{-2}$ |
| G | A | $514 \pm 57.2$ | $0.271 \pm 0.045$ | $5.3 \times 10^{-4}$ | $3.2 \times 10^{-2}$ |
| | T | $476 \pm 112$ | $2.01 \pm 0.44$ | $4.2 \times 10^{-3}$ | $2.6 \times 10^{-1}$ |
| | G | $517 \pm 123$ | $0.125 \pm 0.027$ | $2.4 \times 10^{-4}$ | $1.5 \times 10^{-2}$ |
| | C | $86.1 \pm 11$ | $1.41 \pm 0.13$ | $1.6 \times 10^{-2}$ | 1 |
| C | A | $405 \pm 33.6$ | $1.11 \pm 0.042$ | $2.8 \times 10^{-3}$ | $1.7 \times 10^{-2}$ |
| | T | $479 \pm 80$ | $0.662 \pm 0.045$ | $1.4 \times 10^{-3}$ | $8.7 \times 10^{-3}$ |
| | G | $21.6 \pm 6.3$ | $3.42 \pm 0.35$ | $1.6 \times 10^{-1}$ | 1 |
| | C | $233 \pm 54.2$ | $0.461 \pm 0.05$ | $2.0 \times 10^{-3}$ | $1.2 \times 10^{-2}$ |

Extension efficiency was examined with dGTP, the next correct nucleotide.

To better illustrate the properties of Dpo2 in the nucleotide polymerization, its *f*_inc_ values (x-axis) for inserting a wrong nucleotide opposite a template (Table 1) were plotted against the corresponding *f*^0^_ext_ values (y-axis) for extending from that mispair (Table 2). As shown in Fig. 2, data points are scattered at the upper left part of the figure. These data indicated that Dpo2 exhibits a much higher efficiency in the mismatch extension than in the mispair formation (10^−1^-10^−3^ vs 10^−4^-10^−5^), and the enzyme preferably extends primer termini ended with dG, dA and wobble base-pair (T:G and G:T). These results are in strict contrast to an almost equal efficiency in the mispair formation and in the mismatch extension for Dpo4 and other non-extender DNA polymerases (Johnson et al., 2000; Pavlov et al., 2006), whose data points scatter along the dash line in Fig. S3. Thus, we reasoned that Dpo2 could function as a mismatch extender in *Sulfolobus*.

**Figure 2.**
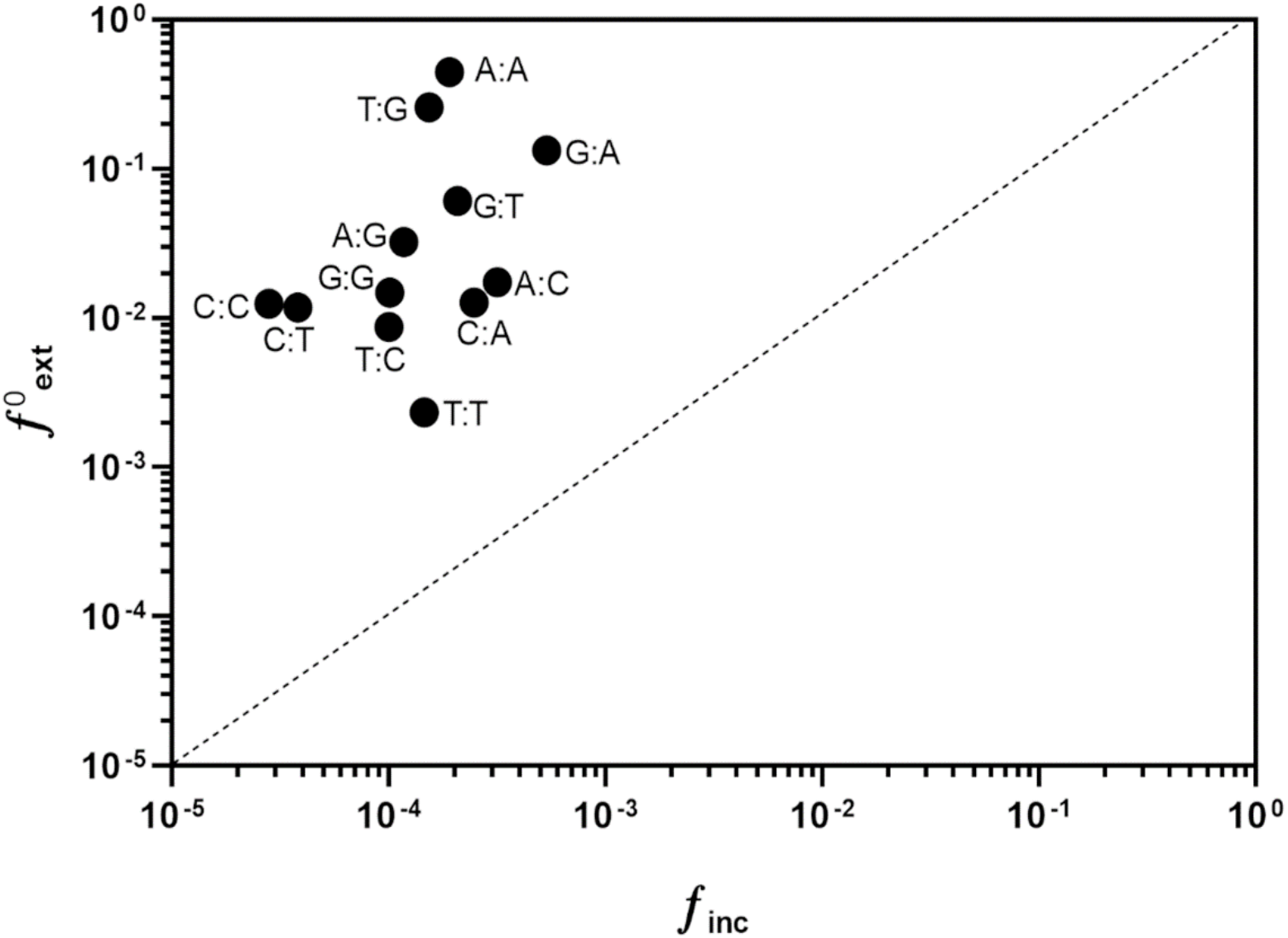
Dpo2 is proficient in extension of mismatched primer termini. Values of *f*^0^_ext_ (the ratio of the apparent K_cat_/K_m_ of extension from the mismatched base pair to the apparent K_cat_/K_m_ of extension from matched base pair) presented in Supplementary Table S4 were plotted against the values of misincorporation frequency (*f*_inc_) shown in Supplementary Table S3. The dash line corresponds to *f*^0^_ext_ = *f*_inc_.

### Dpo2 functions as a DNA lesion extender

The exceptional capability of mispair extension by Dpo2 prompted us to test its activity in translesion DNA synthesis. Three DNA lesions were chosen for the experiment, including AP site, cis-syn cyclobutane pyrimidine dimer (CPD) and 8-oxo-7,8-dihydro-2’-deoxyguanosine (8-oxodG), all of which are common forms of DNA damage encountered by a thermophilic acidophile. The DNA template designed for the AP site bypass experiments was a 37 nt oligonucleotide containing a synthetic abasic site (tetrahydrofuran analogue) at 17th position (Table S2). Two primers were then designed, one extends to the −1 position of the AP site of the template (for “AP insertion” assay), while the other, to the position opposite the abasic site (for “AP extension” assay). Annealing of the DNA template with each of the primers yielded two series of primer-template substrates, for the experiments of “AP insertion” and “AP extension”, respectively. Seven reactions were setup for each assay, in which only the Dpo2 content varied within the indicated range. After incubation for 10 min, primer extension products were analyzed by the denaturing polyacrylamide gel electrophoresis (PAGE). As shown in Fig. 3B, in the AP insertion assay, the signal of primer extension products was hardly detectable at the position across the lesion and beyond even in the presence of 800 nM Dpo2. This enzyme concentration is 100-fold higher than the efficient primer extension on an undamaged DNA template by the polymerase (Fig. 3A). Intriguingly, when the abasic site was covered by the terminal nucleotide of a primer (AP extension), +1 extension product was already detected with 25 nM Dpo2, the lowest enzyme concentration tested here. Furthermore, the size of extension products increased along with the elevation of the Dpo2 concentration, and all primers were converted into longer products at 200 nM enzyme (Fig. 3A). These data indicated that Dpo2 works as an extender polymerase in the TLS bypass of abasic sites.

**Figure 3.**
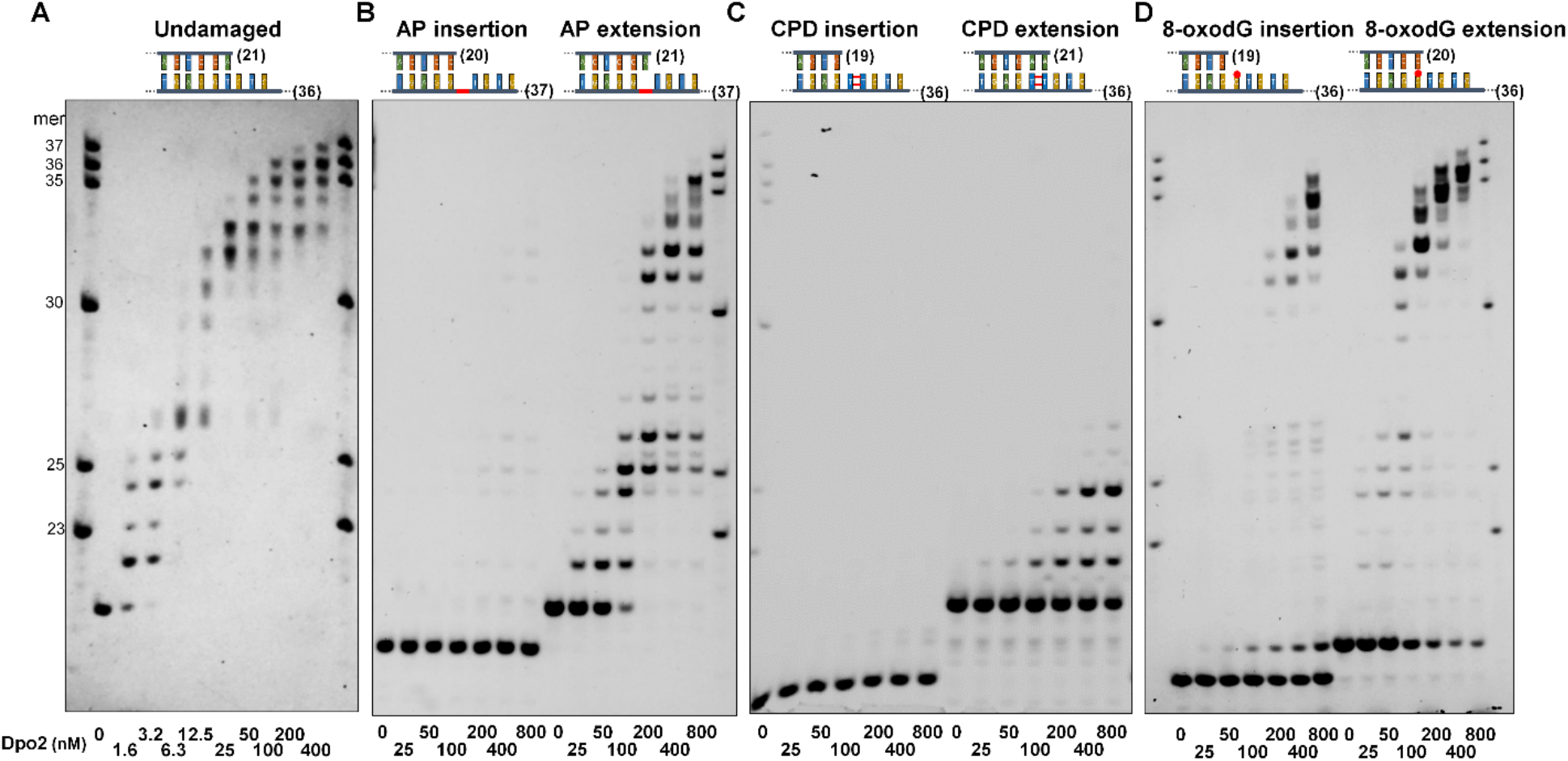
Dpo2 is a lesion extender. DNA substrates employed for primer extension assay were illustrated above the corresponding gel images. Templates in the substrates are of 4 different types: (A) Undamaged template, which is either lesion-free (undamaged); (B) template carrying an AP lesion, which is highlighted in red in the backbone; (C) template carrying TT-CPD, which is shown as two parallel bars adjoined with two red lines, and (D) template containing 8-oxodG (shown as “G” base carrying a red hat). Numbers in parentheses indicate lengths of primers and templates in each substrate. Primer extension was conducted with reaction mixes containing Dpo2 of varied concentrations (indicated below gel images) and analyzed by denaturing PAGE. Numbers in the size marker denote the lengths of nucleotides.

When a DNA template carrying a cis-syn cyclobutane pyrimidine dimer (CPD) was employed, we found again that Dpo2 failed to insert any nucleotide opposite the lesion, but it was capable of extending mispaired primer ends, albeit at an efficiency lower than the extension of the AP-contained mispaired primer ends (Fig. 3C). In the case of the 8-oxodG bypass reaction, Dpo2 incorporated a nucleotide across the 8-oxodG lesion for 53.8% of the template even at the highest concentration (800 nM) of the enzyme tested in this study, indicative of a very weak activity in the TLS insertion. In contrast, the Dpo2 extension is robust since a comparable amount of substrate has rapidly been extended at an 8-fold lower concentration (100 nM) (Fig. 3D).

### *S. solfataricus* Dpo2 is also a robust DNA polymerase lacking proofreading

We noticed that our results with the *S. islandicus* Dpo2 are in contrast to those obtained with the *S. solfataricus* Dpo2 (SsoDpo2) that expressed in an *E. coli* host in a previous work. In the latter only weak activities were observed in polymerization and in proofreading for the heterlogously expressed form of SsoDpo2 [37]. Since Dpo2 proteins of *S. solfataricus* and *S. islandicus* share 91%/96% sequence identity/similarity, it is very unlikely the two proteins would exhibit any major differences in enzymatic properties. SsoDpo2 was then expressed in *S. islandicus*, and the enzyme was purified and characterized along with the *S. islandicus* Dpo2 (Fig. S5). We found that, as for the *S. islandicus* Dpo2, the recombinant SsoDpo2 obtained from the *Sulfolobus* host showed two striking features: (a) it does not possess any detectable 5’-3’ exonuclease activity, which is consistent with the lack of the Exo motifs, and (b) the PolB enzyme is highly efficient in nucleotide incorporation, exhibiting a strong propensity in mismatch extension and AP extension during DNA synthesis (Fig. 4).

**Figure 4.**
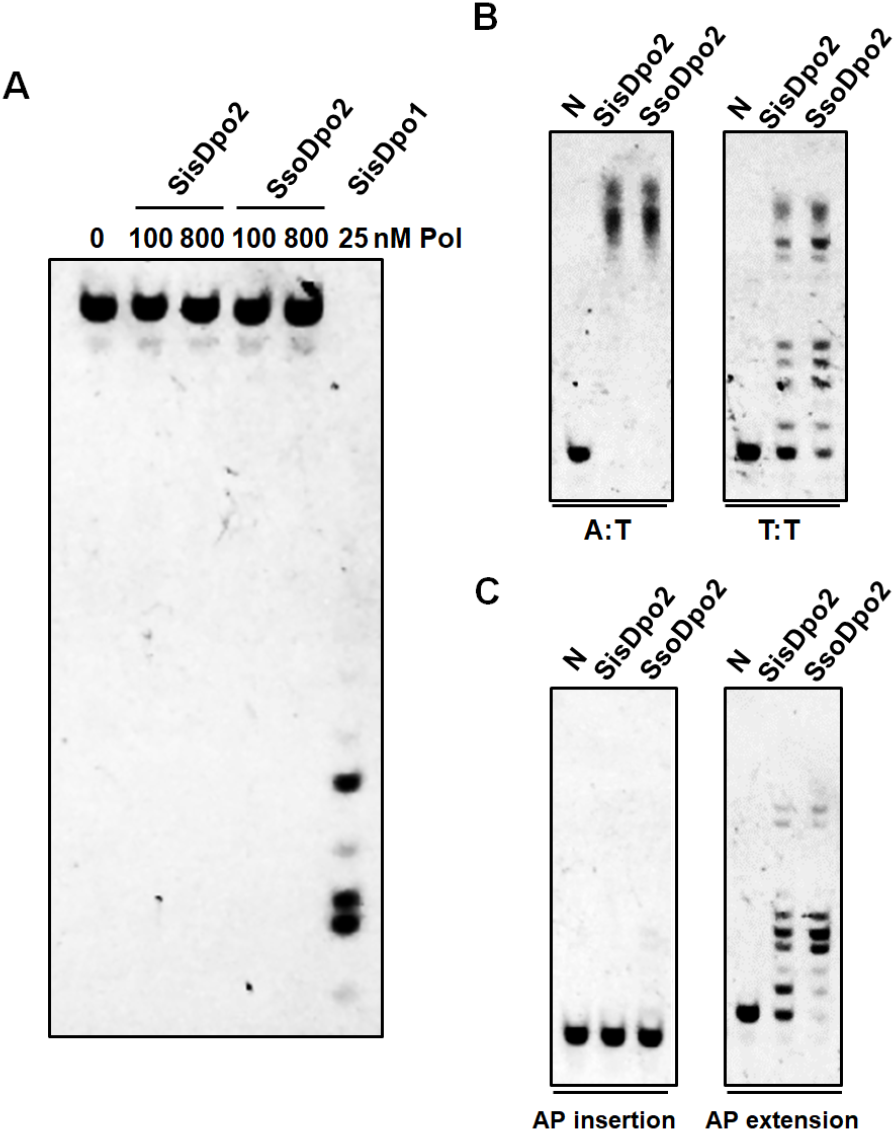
SsoDpo2 has a similar activity as SisDpo2. (A) Exonuclease assay. The assay was set up with 50 nM substrate and enzyme concentrations indicated above each lane. Reactions were conducted at 60 °C for 5 min. (B) Extension of the undamaged substrate (A:T) and mismatched substrate (T:T). Substrates used for the assays are the same as shown in Fig. 1. Each reaction contains 100 nM DNA polymerase and 50 nM substrate. N: no enzyme control. (C) AP insertion and extension. Assays were set up with the substrates shown in Fig. 3B. Primer extension reactions were conducted with 50 nM substrates.

Taken together, the above results indicated that PolB2s are very inefficient in the misincoporation and TLS insertion, but show robust activity in mismatch and lesion extension. Thus they probably function as mismatch and lesion extenders in the archaeal translesion DNA synthesis.

## DISCUSSION

Members of PolB2 enzymes are widespread in Archaea. This group of DNA polymerases are unique since they carry deletions or radical variation in exo motifs and variations in key amino acid residues of the PolC motif that are very conserved in all other groups of PolB enzymes [21; 38]. For this reason, PolB2 was regarded as a group of inactivated DNA polymerases. Here, we report that both *S. islandicus* and *S. solfataricus* Dpo2 enzymes are very active in nucleotide incorporation since their activities are comparable to that of Dpo1, the replicase that co-exists in these archaeal organisms. We further show that the archaeal PolB2s lack any detectable 5’-3’ nuclease activity and they are proficient in mismatch and lesion extension. Our results suggest the archaeal PolB2 enzymes represent a novel type of PolB that play important roles in DNA damage repair.

Our identification of the robust DNA polymerase activity for the *Sulfolobus* Dpo2 has yielded important insights into the mechanisms of DNA synthesis. B-family DNA polymerases share conserved motifs three of which (Exo I, II, III) are located in the proof-reading domain while the remaining (e.g., PolA, B, C) are in the polymerase domain (Fig. 1A). The PolB2 group of DNA pols exhibit numerous variations in these motifs. The *Sulfolobus* Dpo2 proteins investigated in this work represent the smallest PolB2 known to date (Fig. S6). These enzymes lack most of the conserved amino acids in the proofreading domain and exhibit large variations in the PolC motif. The latter is in contrast to the members of other PolB groups since their PolCs have the YxDTD invariant motif, which is replaced with HxxxD in PolB2. It has been reported that PolC motif of B-family DNA pols plays a key role in primer/template recognition and participates in the coordination of the catalytic Mg^2+^ that is essential for the polymerization reaction [39; 40]. Nevertheless, previous works have already shown that the two Asp residues in PolC are not equally important for catalysis of DNA polymerization. Structural interrogation of a few B-family DNA polymerases has revealed that the second aspartate is responsible for the metal ion coordination whereas the first Asp is orientated away from the activity center [40; 41; 42; 43; 44]. Since mutagenesis of the first Asp greatly reduces the activity of the human Pol a and two viral replicases [45; 46; 47], this acidic amino acid, although not directly involved in catalysis, still plays an important role in the polymerase activity. However, our work shows that *Sulfolobus* Dpo2 enzymes, although lacking the first Asp of PolC, are as active as the Dpo1 replicase in nucleotide incorporation. This suggests the function of the first Asp in the PolC motif can be functionally replaced by His, an invariant amino acid in the PolC motif of the PolB2 enzymes (Fig. 1A). In addition, mutation of amino acid residues adjacent to the catalytic Asp in the PolC motif of the *E. coli* Pol I or *Thermus aquaticus* (Taq) polymerase impairs their mismatch extension ability [48; 49]. To this end, we reason that the sequence variation at the PolC motif in the PolB2 enzymes could reflect their adaptation to their specialized function in DNA repair, which apparently requires a robust nucleotide incorporation activity and the tolerance of DNA damage whereas their replication processivity and proofreading are disfavored.

Noticeably, the properties of the recombinant SsoDpo2 we have obtained from *S. islandicus*, a homologous host are very different from the same enzyme yielded from heterologous expression in *E. coli* [37]. While the *E. coli* recombinant SsoDpo2 (500 nM) exhibits optimal activity at 50 °C, and the activity is greatly reduced at 60 °C and completely inactivated at 70 °C [37], the optimal temperature for the *Sulfolobus-expressed* SsoDpo2 (17.5 nM) is 50-65 °C and the enzyme is still active in 80-90 °C (Fig. S5). We reason that the observed differences can be attributed to differences in posttranslational modifications (PTMs) present in proteins produced in the thermophilic host versus those synthesized in the mesophilic host [50]. Indeed, in a comparative study of a recombinant *S. islandicus* esterase produced in *S. islandicus* versus that produced in *E. coli*, the homologously expressed protein is much more active than the heterologously expressed version of the same enzyme [51]. In addition, Dpo2 contains 7 cysteine residues and their potential of generating intra- and/or intermolecular disulfide bonds may differ strongly in a different genetic background, which also contribute to the differences observed between the two forms of SsoDpo2 recombinant protein.

Our characterization of the archaeal Dpo2 enzymes has revealed that PolB2 enzymes exhibit several distinctive biochemical features, including: (a) the lack of a proof-reading activity, (b) its promiscuous extension of mispaired primer ends can fix mismatches and (c) its capacity in lesion extension may generate mutations. The unique features are consistent with their possible functions in DNA damage repair in these crenarchaea as recently revealed from our genetic analyses in *S. islandicus* using the gene disruptant strains for *dpo2, dpo3*, and *dpo4*, respectively. Comparison of their phenotypes with that of the wild-type reference has revealed that Dpo2 is solely responsible for the targeted mutagenesis in this crenarchaeon [20]. These unique features of Dpo2 may have provided the molecular mechanisms for the generation of the Dpo2-dependent targeted mutagenesis observed in our genetic study [20]. In this regard, Dpo2 is analogous to the eukaryotic Pol ζ since its deficiency also reduces the targeted mutations in yeast [52; 53] and this B-family DNA polymerase also known for the lack of proofreading activity and the exceptional ability in mismatch extension [54]. Considering Pol ζ working in concert with Y-family DNA polymerases (Pol ι or Pol η) in a two-polymerase mechanism for AP lesion bypass in Eukarya [29; 55], Thus, the identification of PolB2 enzymes as a mismatch and lesion extender raises an intriguing question about whether it also acts in concert with other DNA polymerase in Archaea to facilitates lesion bypass in the domain of Archaea.

## AUTHOR CONTRIBUTIONS

QS and XF designed the work. XF, BZ, ZG, RX, XL, MF, SI, YS, YI and QS contributed to the acquisition and analysis of the data. XF, BZ, QS, SI, YI and YS interpreted the data. QS and XF wrote the manuscript, and SI, YI and YS revised it.

## ACKNOWLEDGEMENT

We thank Dr. Likui Zhang at Yangzhou University for advices in kinetic studies and Dr. Li Huang at Institute of Microbiology, Chinese Academy of Sciences for stimulating discussions.

## DATA AVAILABILITY

All data required to evaluate the conclusions of this study can be found in either the main text or the Supplementary data.

## Supplementary Material

### Supplementary Tables

**Supplementary Table S1.**
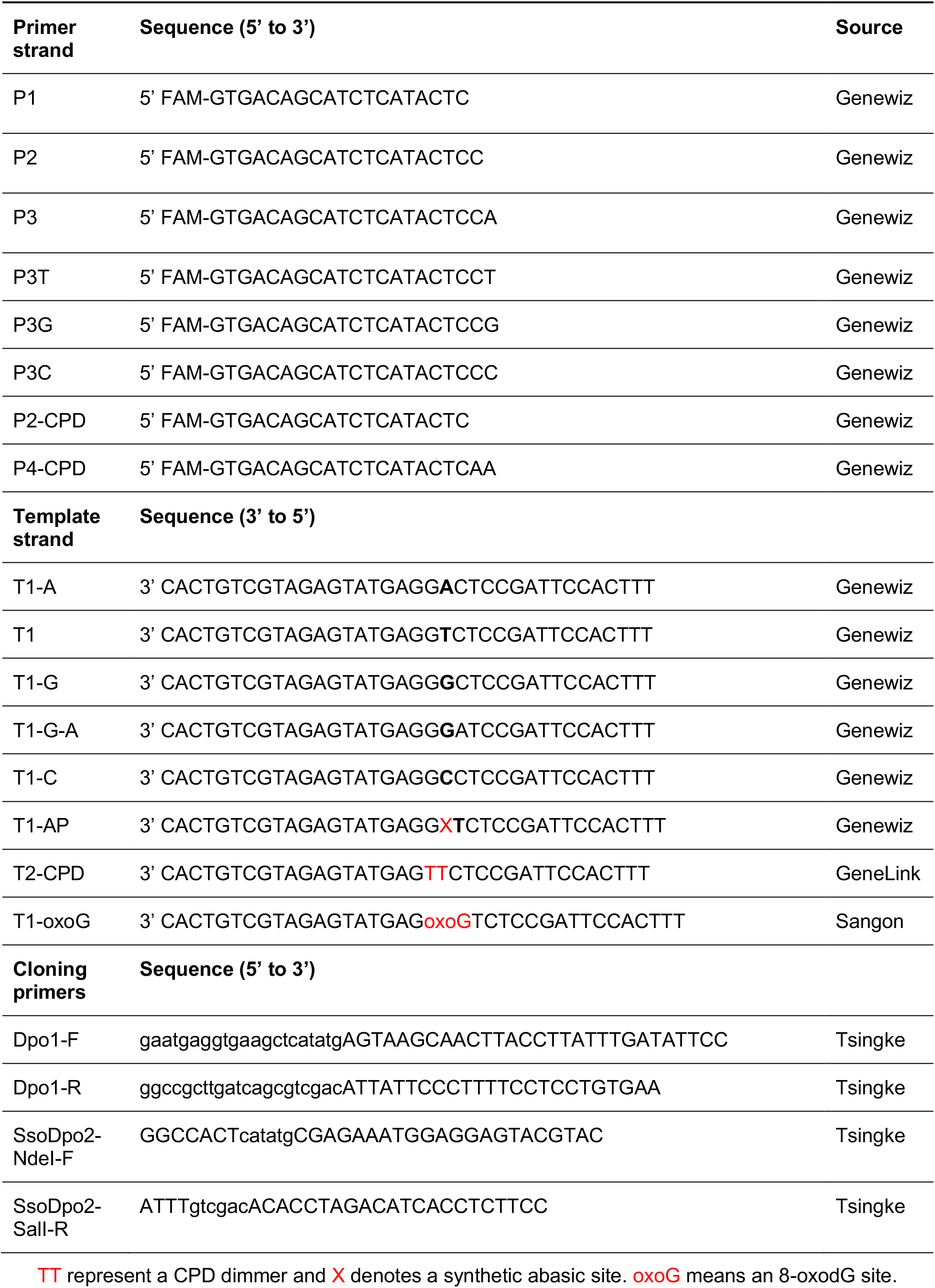
Oligonucleotides Used in This Study.

**Supplementary Table S2.**
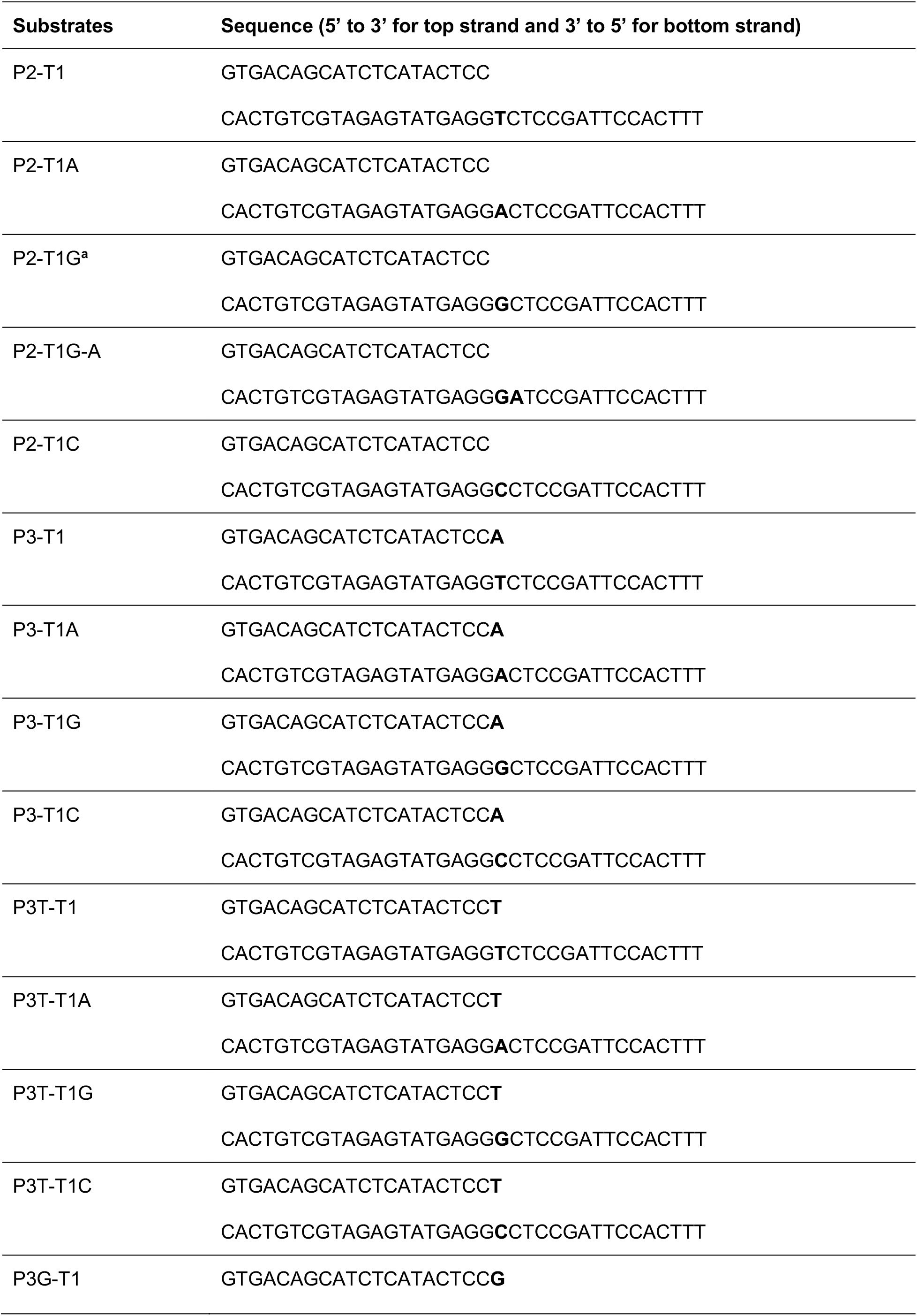
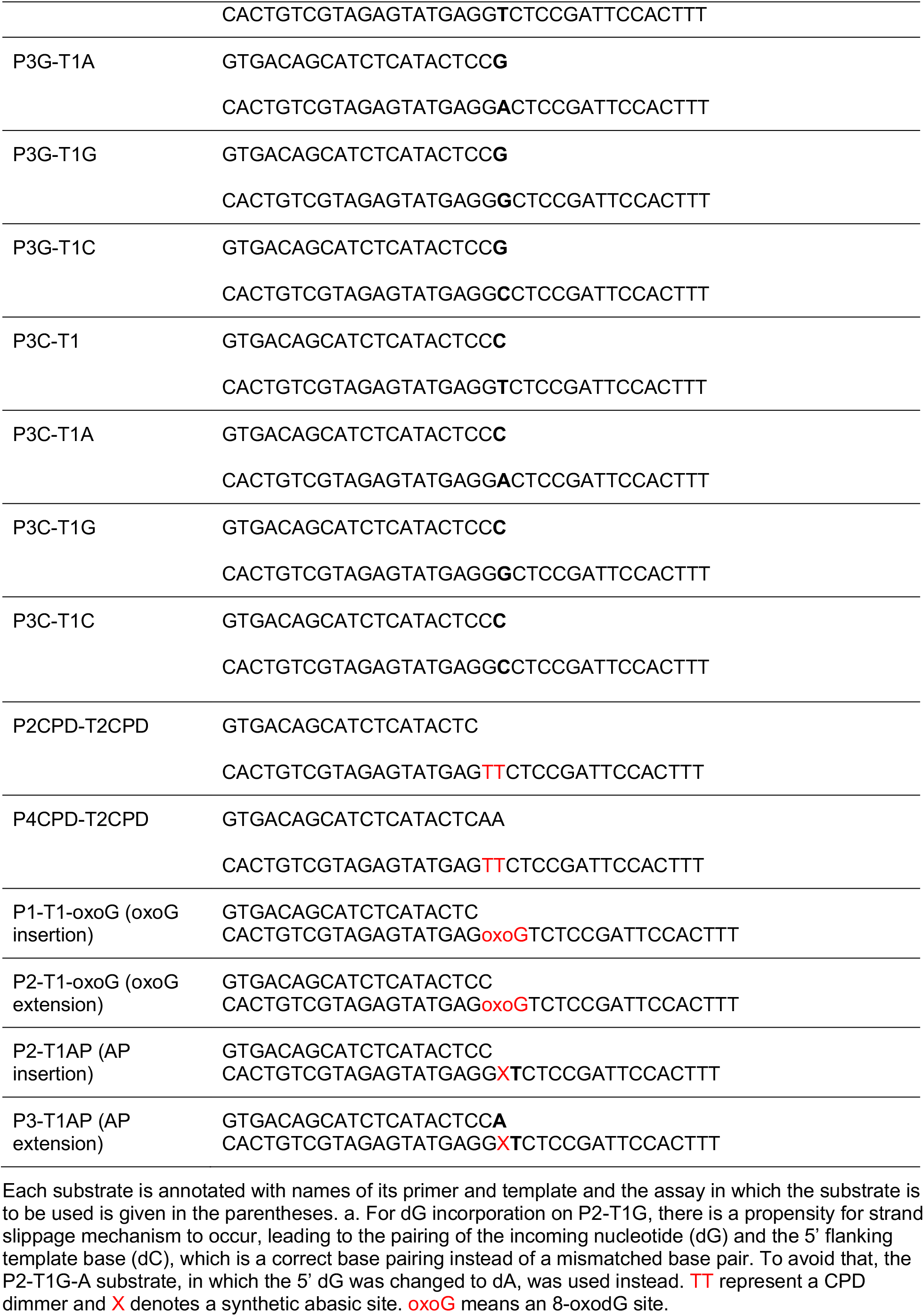
DNA Substrates Used in This Study.

**Supplementary Figure S1.**
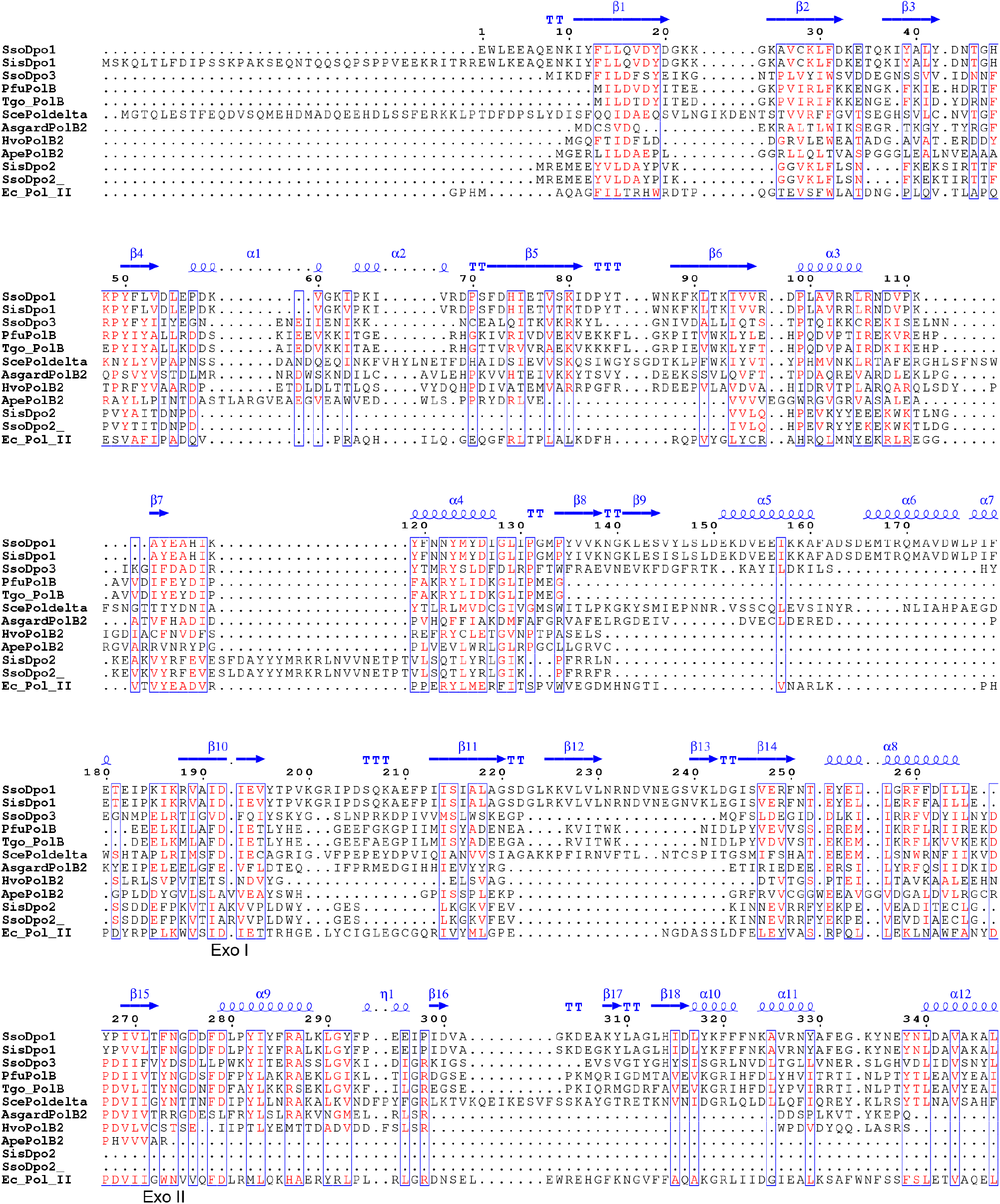
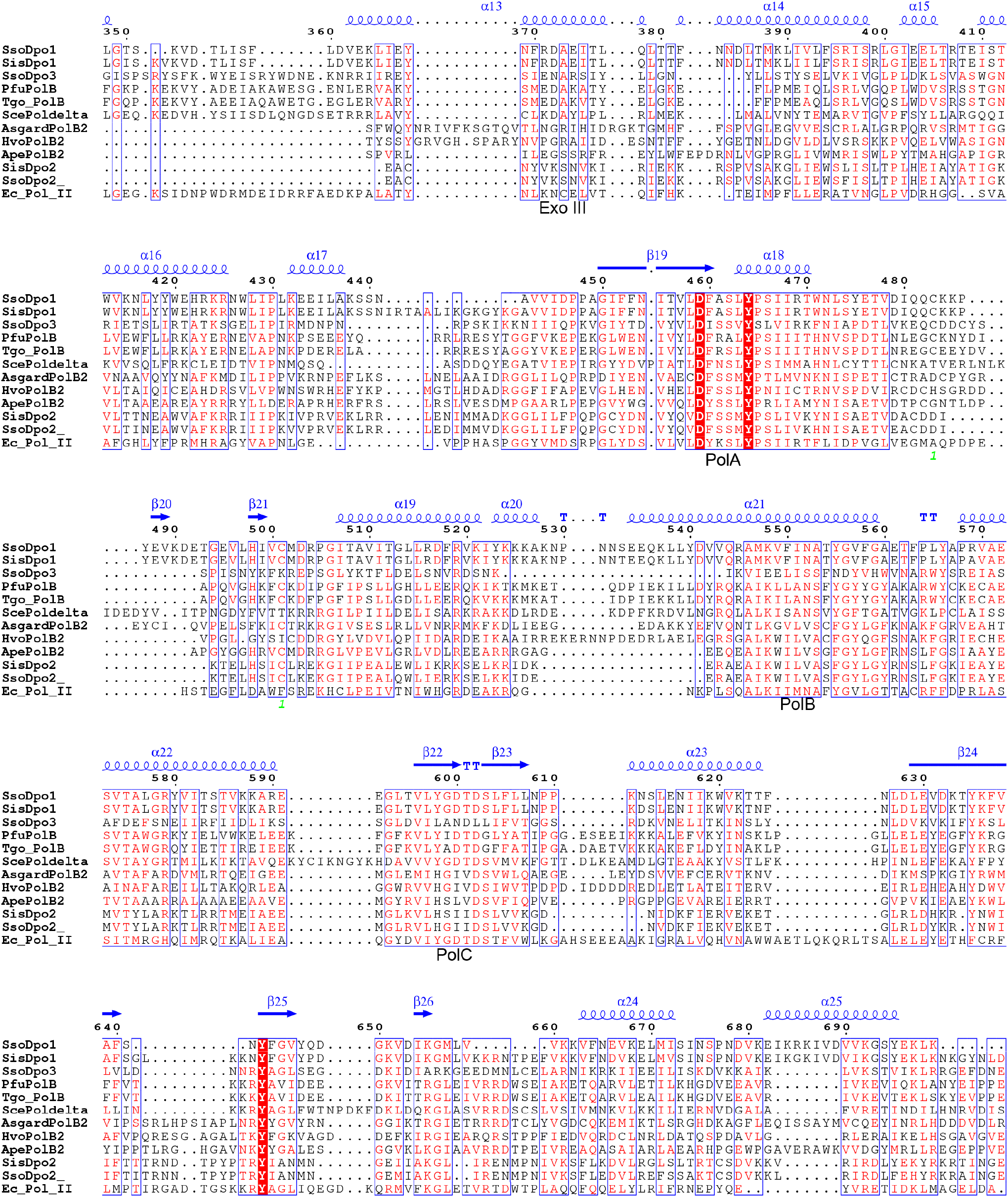
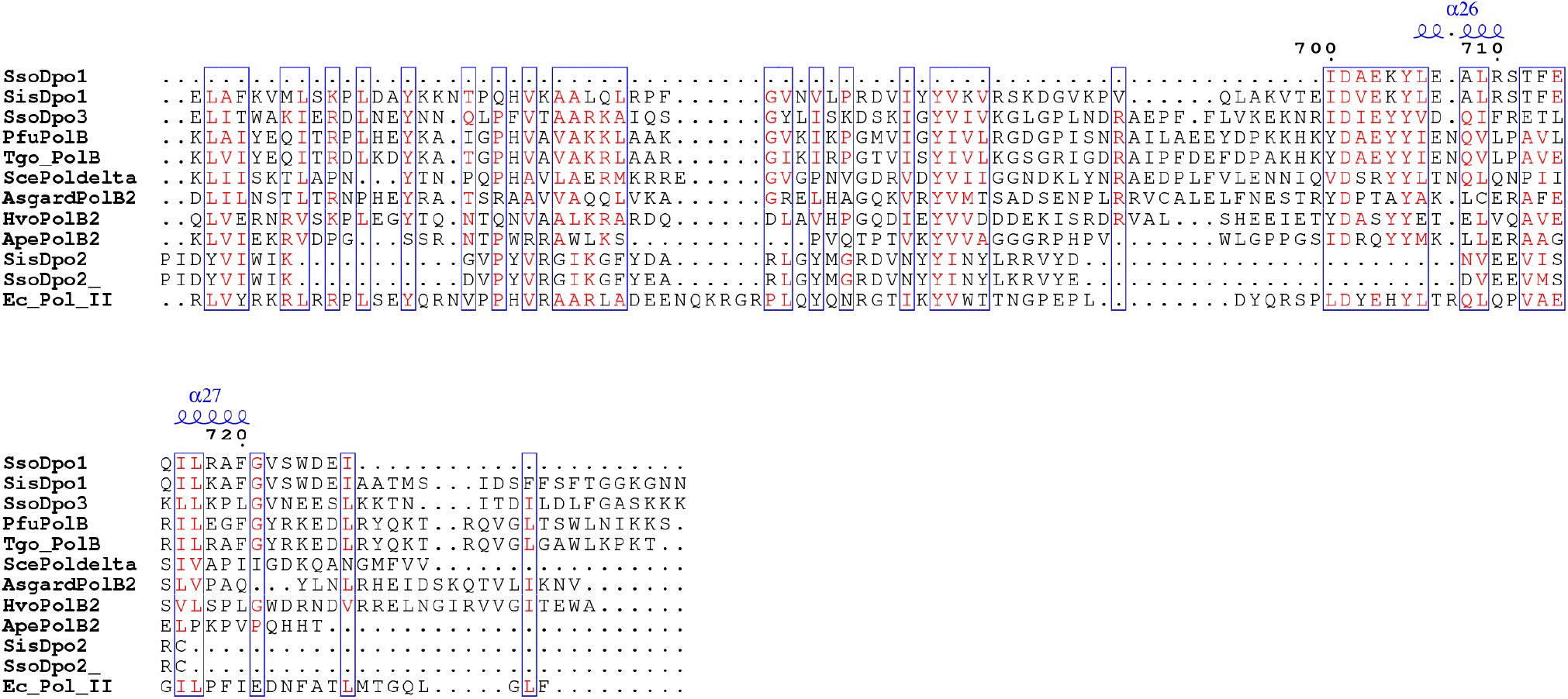
Sequence aligment of a selelcted set of B-family DNA polymerases. B-family DNA Polymerases employed for the analysis include SsoDpo1, *S. solfataricus* Dpo1; SisDpo1, *S. islandicus* Dpo1; SsoDpo3, *S. solfataricus* Dpo3; PfuPolB, *Pyrococcus furiosus* PolB; Tgo_PolB, *Thermococcus gorgonarius* PolB; ScePoldelta, the catalytic subunit of *Saccharomyces cerevisiae* Pol δ; AsgardPo1B2, Candidatus Thorarchaeota archaeon PolB2; HvoPolB2, *Haloferax volcanii* PolB2; ApePolB2, *Aeropyrum pernix* PolB2, and Ec_Pol_II, *E. coli* Pol II. Structures of SsoDpo1 (1S5J) was used as the templates for the structure-based sequence alignment. The secondary structural elements shown above the sequences were retrieved from the structure file of SsoDpo1 (1S5J).

**Supplementary Figure S2.**
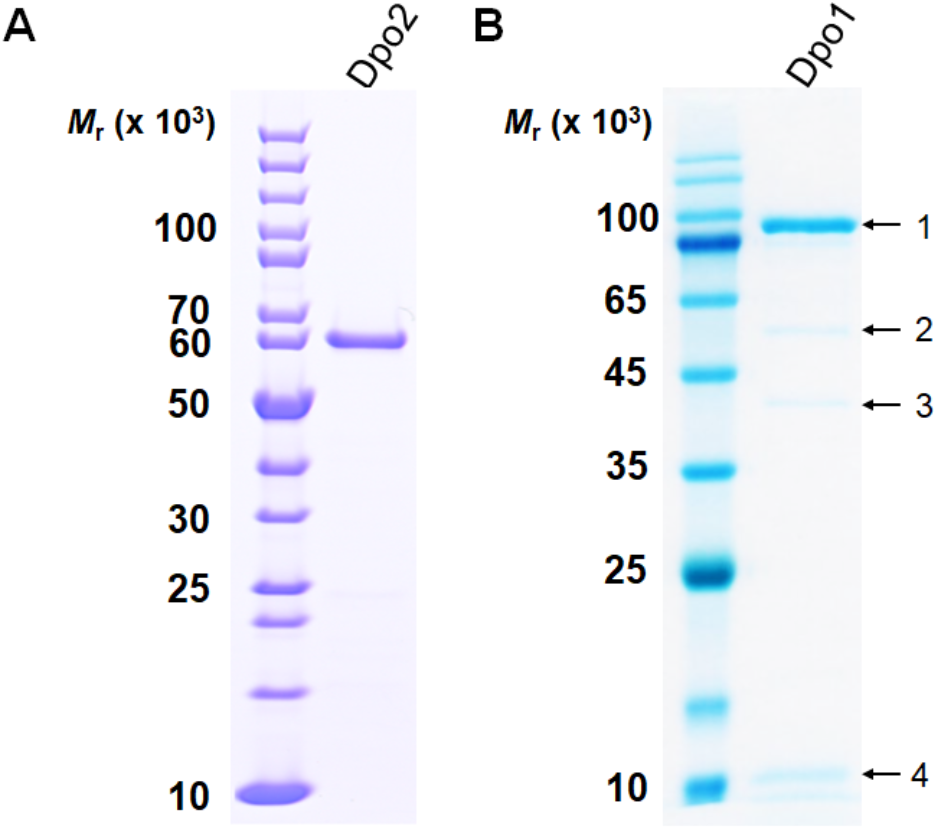
SDS-PAGE analysis of purified Dpo2 and Dpo1 proteins from the native host. (A) Dpo2 with a theoretical size of 64,927 daltons. (B) Dpo1 with a theoretical size of 101,205 daltons. Mass spectrometry confirmed that the band 1 corresponsding to a size of 100,000 was PolB1, as were two smaller species of ~60,000 (2) and ~40,000 (3), which were presumably resulted from Dpo1 degradation. The smallest species (4) were identified as Dpo1 degredation and PBP1 (SiRe_1861), which is probably part of the Dpo1 holoenzyme, as have been shown for SsoDpo1 [8].

**Supplementary Figure S3.**
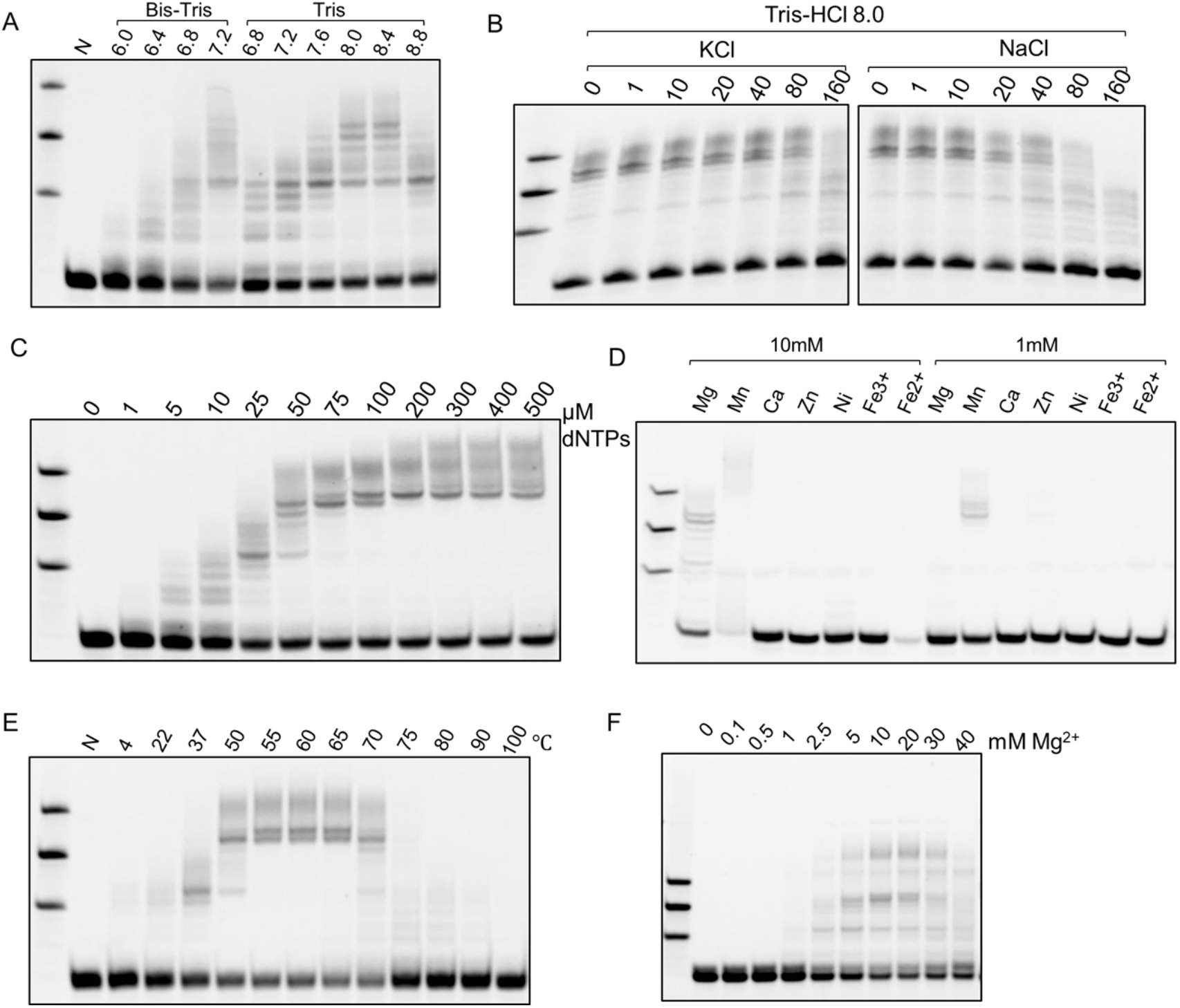
Optimization of the reaction condition. DNA substrates were the same as shown in Figure 1. Reactions were set up with 37.5 nM Dpo2 and 50 nM substrates. (A) Optimization of the reaction pH at 60 °C. N: no enzyme control. (B) Optimization of salt concentration at 60 °C using the Tris-HCl buffer, pH 8.0. (C) Optimization of dNTP concentration, using Tris-HCl pH 8.0 buffer containing 40 mM KCl and 0.1 mg/ml BSA at 60 °C. (D) Metal ions-dependence of Dpo2. The reactions were carried out with Tris-HCl pH 8.0 containing 40 mM KCl and 0.1 mg/ml BSA at 60 °C. Mg^2+^ was used since some of the following analyses involved the use of multiple DNA Pol enzymes and a much higher physiological concentration of Mg2+ have been reported than other metal ions. (E) Determination of theoptimal reaction temperature using Tris-HCl pH 8.0 buffer containing 40 mM KCl and 0.1 mg/ml BSA at indicated temperature. (F) Optimization of Mg2+ concentration. At last, the optimized buffer system for Dpo2 contains 50 mM Tris-HCl pH 8.0, 40 mM KCl, 0.1 mg/ml BSA, 10 mM MgCl_2_, 100 μM dNTPs. In the single nucleotide incorporation assay and kinetics analysis, the dNTPs mixtures were omitted and single dNTP with indicated concentrations was instead used.

**Supplementary Figure S4.**
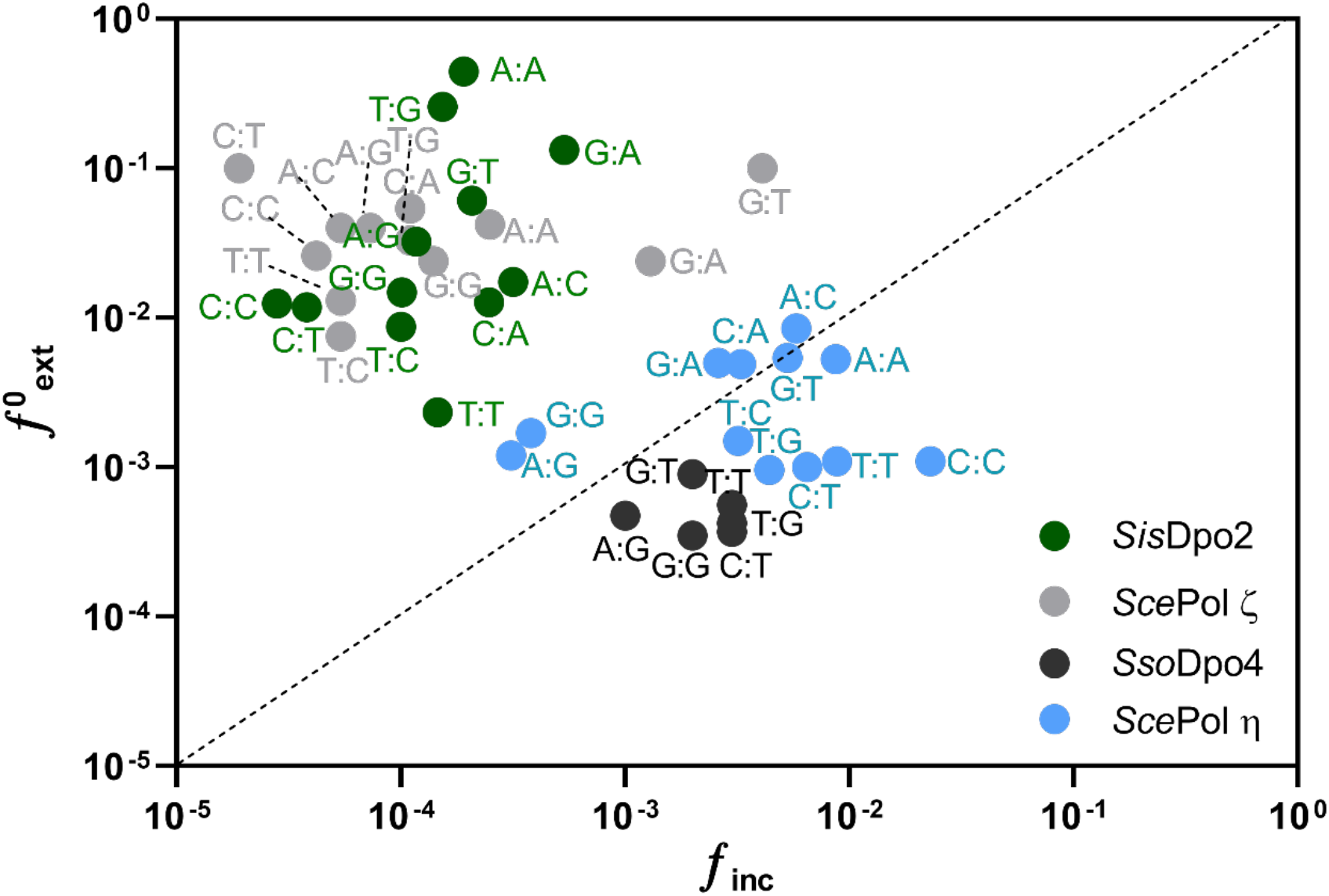
A comparison of mismatch extension ability of extender and nonextender DNA polymerases. The *f*inc and *f*0ext values of Saccharolobus solfataricus Dpo4 (SsoDpo4), Saccharomyces cerevisiae Pol ζ (ScePol ζ) and S. cerevisiae Pol η (ScePol η) are retrieved from [56], [29] and [57] respectively.

**Supplementary Figure S5.**
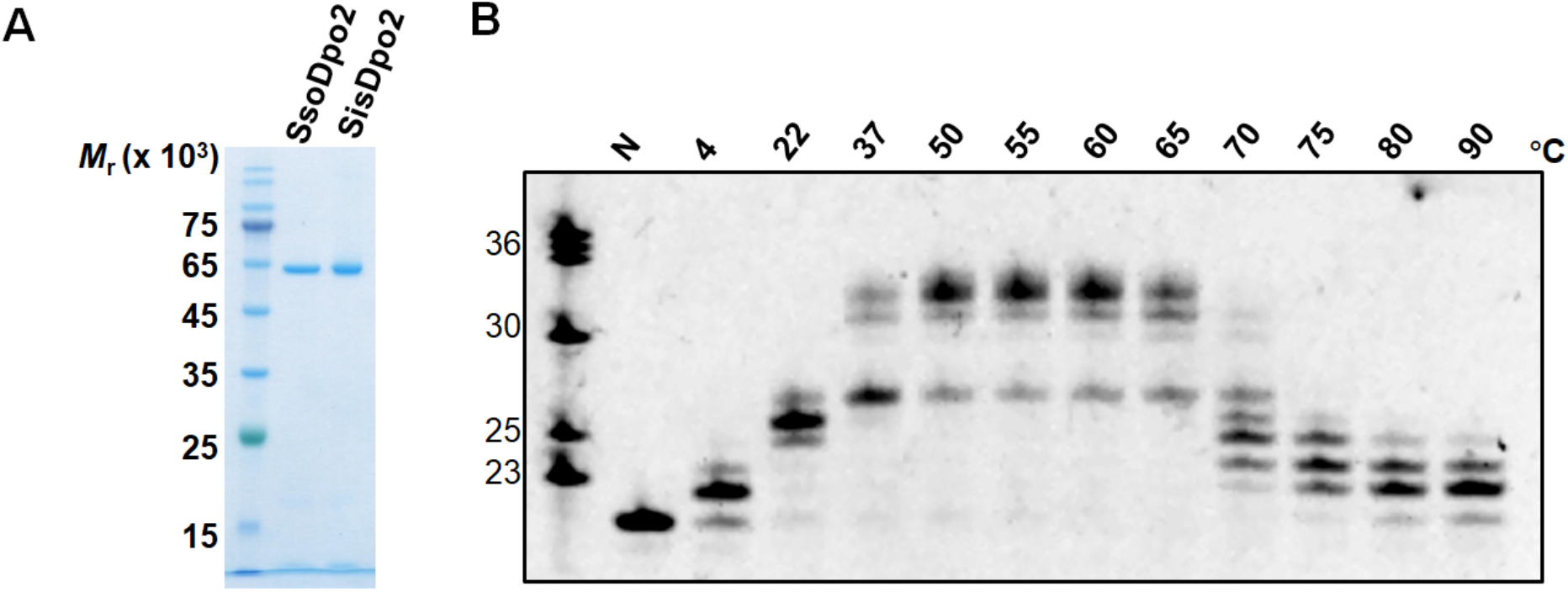
the optimal temperature of SsoDpo2. (A) SDS-PAGE analysis of purified SisDpo2 and SsoDpo2 proteins. SisDpo2, Sulfolobus islandicus Dpo2. SsoDpo2, *S. solfataricus* Dpo2. Both proteins were expressed in the S. islandicus host. (B) the optimal temperature of SsoDpo2. The reaction contains 17.5 nM SsoDpo2 protein and 50 nM substrate used in Fig 1B, in the presence of 100 μM dNTPs. The experiment was carried out at the temperature indicated above each lane for 5 min.

**Supplementary Figure S6.**
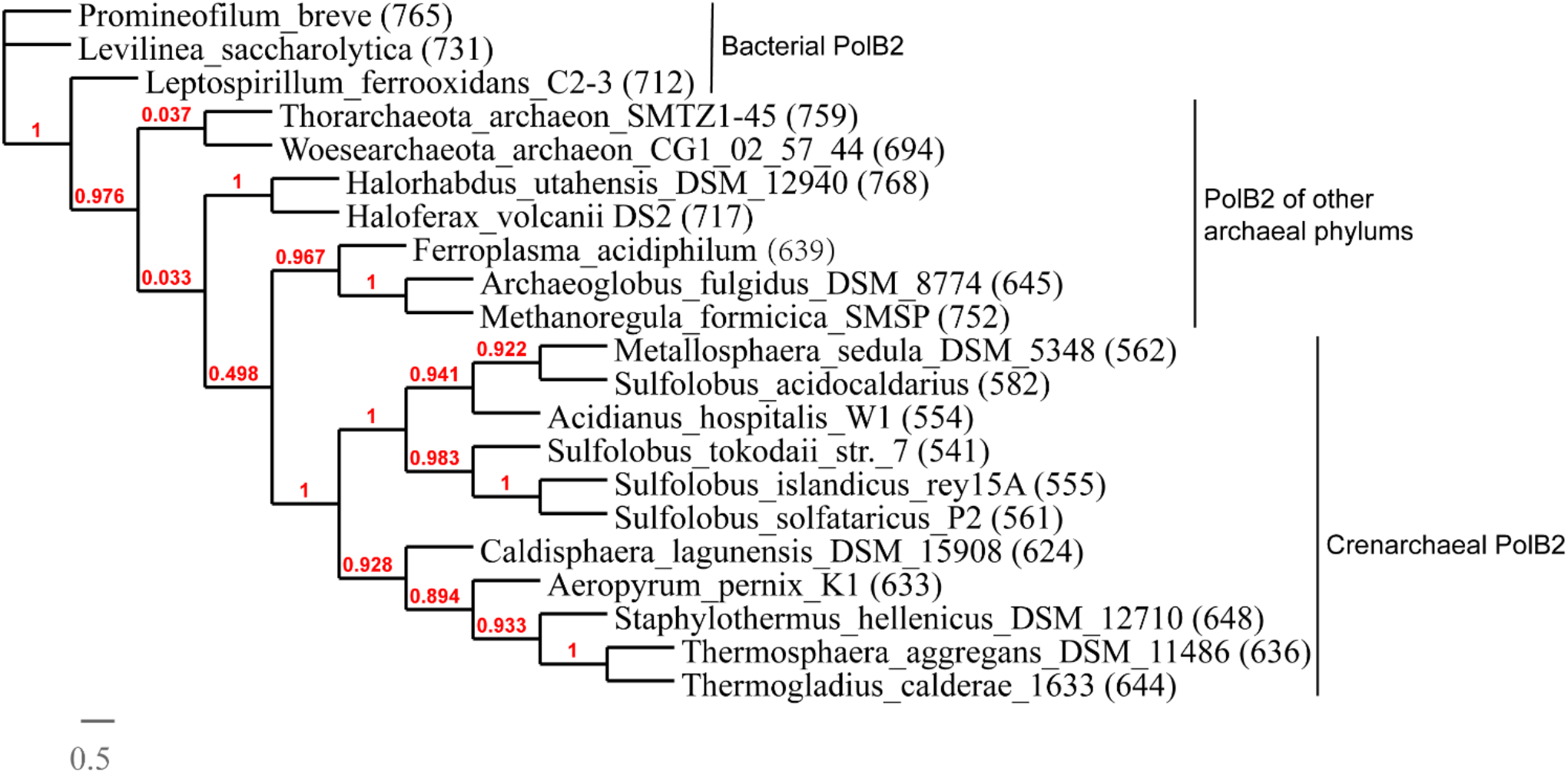
The phylogeny of PolB2 proteins. The phylogenetic tree was constructed using sequences of PolB2s extracted from NCBI and the analysis was performed on the Phylogeny.fr platform [58]. The size of each PolB2 was indicated by the numbers in parentheses (aa).

